# Empirical evidence of resource dependent evolution of payoff matrices in *Saccharomyces cerevisiae* populations

**DOI:** 10.1101/2024.01.15.575778

**Authors:** Pavithra Venkataraman, Anjali Mahilkar, Namratha Raj, Supreet Saini

## Abstract

In evolutionary game theory, a relative comparison of the cost and benefit associated with obtaining a resource, called payoff, is used as an indicator of fitness of an organism. Such payoff matrices are used to understand complex inter-species and intra-species interactions like cooperation, mutualism, and altruism. In the absence of any empirical data, the evolution of these payoff matrices has been investigated theoretically by tweaking well-established game theory models. In this paper, we present empirical evidence of three types of resource-dependent changes in the payoff matrices of evolving *Saccharomyces cerevisiae* populations. We show that depending on the carbon source and participating genotypes, the payoff matrix could either (a) evolve quantitatively yet maintain a cheater-cooperator game, (b) change qualitatively such that the cheater-cooperator game collapses, or (c) change qualitatively to result in the birth of a cheater-cooperator game. Our results highlight the need to consider the dynamic nature of payoff matrices while making even short-term predictions about population interactions and dynamics.

## Introduction

Impalas living in the Savannahs groom themselves with their teeth, tongue, and saliva to get rid of harmful ticks from their bodies. However, since an impala cannot reach all parts of its own body, it needs other members of the population to groom it^1^. Since grooming another individual comes with costs, every impala in the population has a choice – should it groom the other members (cooperate), or not (defect)? Such “games” are ubiquitous across life forms, and make for examples of Prisoner’s dilemma, a classic game theory problem.

A payoff (or fitness) matrix associated with the Prisoner’s Dilemma game is as shown below.

|  |  | <u>Opponent’s move</u> |  |
| --- | --- | --- | --- |
|  |  | Cooperate | Defect |
| Cooperate |  | 3 | 0 |
| Defect |  | 5 | 1 |

The structure of the payoff matrix dictates that irrespective of what the opponent is doing, it always pays more to defect. Thus, the stable equilibrium of the game is to defect, for both players. However, cooperation still survives in nature.

One explanation for the prevalence of cooperation is that repeatedly playing the game (an impala needs grooming from its herd throughout its life) ensures that the strategy to “defect” would be found out soon^2^. In such a context, explanation of existence of cooperation as a natural phenomenon becomes feasible.

In recent years, game theory has been used to study interactions among members of a microbial population^3–8^. Microbial systems studied in this context involve a “public good” – which could be a siderophore, toxin, or an enzyme. A cell secreting the “public good” pays for the cost of production of the good, while the benefit of the good is reaped by all members of the population. In this scenario, cooperators utilize several mechanisms to ensure that the cheater population does not drive them to extinction^9^.

While studies shed light on how microbial interactions could be looked at through the prism of game theory^10^, they are limited by their inability to predict how the payoff matrix itself evolves with time. Specifically, given large population sizes, members of the participating populations will accrue beneficial mutations, and how these mutations change the rules of the “game” and consequently, the payoff matrix is not known.

How could commitment to cooperation, and consequently, the payoff matrix change?^11^ What is the relative abundance of the possibilities? How do specific ecological contexts “force” populations to adopt one strategy over the other? All these questions remain unanswered.

Several theoretical works have addressed this, and proposed possibilities. These include the evolution of a Prisoner’s dilemma to a Hawk-Dove game^12^, the emergence of a stable polymorphism in the system^12–14^, a complete collapse of the system^14^, and increased payoff leading to increased investment in cooperation^15^. In the absence of knowledge regarding the connect between the genotypic changes and the phenotype, the payoff matrix evolution is often simply modelled via random variables^16^.

There is no empirical data to explain evolution of payoff matrices. Understanding these transitions is important not only from the context of enhancing how much we know of microbial interactions but also to understand stability of microbial communities in an ecological context.

To answer these questions, in this work, we report the evolution of payoff matrices in three examples of “public good” systems in the yeast *Saccharomyces cerevisiae* (*S. cerevisiae*). We show that the payoff matrix associated with Prisoner’s dilemma can change drastically and dramatically - changes to the payoff matrix can be quantitative or qualitative. Our empirical results highlight that while game theoretic frameworks can be representations of biological systems, they have limited ability to predict transitions, even in an extremely short timeframe.

## Results

### Case I. Quantitative alterations to the payoff matrix of S. cerevisiae populations in sucrose

Sucrose, a glucose-fructose disaccharide, is hydrolysed in the periplasm of *S. cerevisiae* by an invertase encoded by SUC2 (**Figure 1A**). An overwhelming majority of the resulting monosaccharides diffuse in the surrounding media; while the cell producing SUC2 only retains ∼2% of the monosaccharides^17^. Thus, SUC2 encodes for a public good, whose benefit (monosaccharides) is reaped even by the non-producers. Cells lacking SUC2 are unable to grow in media containing sucrose as the sole carbon source^18^.

**Figure 1.**
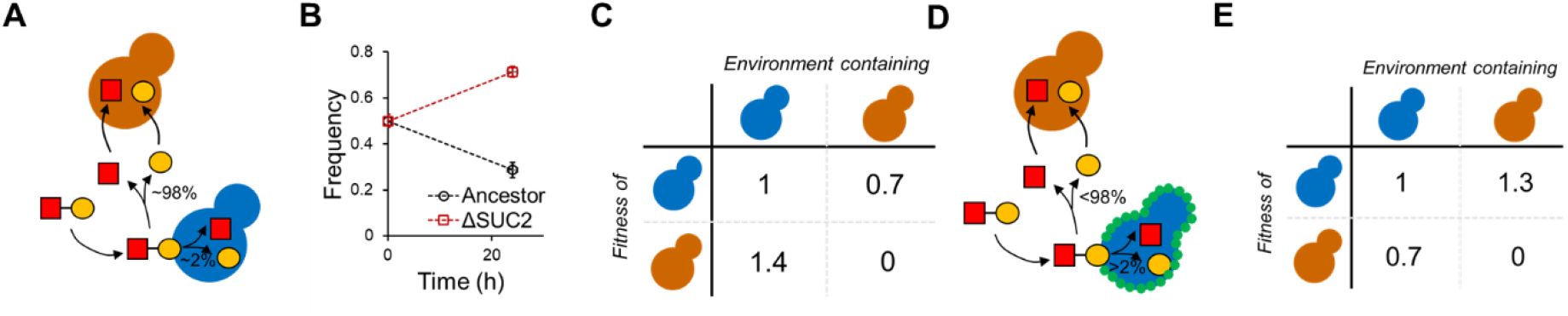
Quantitative changes in the payoff matrix upon co-evolution of a cheater-cooperator pair in sucrose for 200 generations. **(A)** Sucrose hydrolysis takes place in the periplasm of the cooperator (blue). ∼98% of the resulting monosaccharides are released in the media and are available for anyone in the population. The cheater (orange) grows by utilizing the glucose and fructose released by the cooperator. The two monosaccharides are represented by the yellow circle and red square. **(B)** When co-cultured in a 1:1 ratio and allowed to grow for 24 h, the cooperator: cheater ratio decreases to ∼3:7. **(C)** The payoff matrix of the system is as shown when a small number of cheaters/cooperators are introduced in a population of cooperators/cheaters. Cheaters perform better than cooperators, when present in a population of cooperators. **(D)** Upon evolution for 200 generations, a duplication of *HXT* gene likely results in increased privatization of the monosaccharides. The HXTp transporter is indicated via green circles on the cell membrane. **(E)** This duplication results in change in the payoff matrix between the co-evolved cheater and cooperator.

When co-cultured, the cells lacking SUC2 (cheaters) grow to higher frequencies when compared to the cells expressing SUC2 (cooperators) (**Figure 1B**), and the payoff matrix associated with this system is as shown in **Figure 1C**.

However, when the cheater-cooperator pair is co-evolved in sucrose for ∼200 generations, Raj and Saini^19^ report how cooperators in a cheater-cooperator population thrive by increasing resource privatization of the monosaccharides, in this environment. Resource privatization is achieved by the duplication of a high affinity glucose transporter in the cooperator (**Figure 1D**).

As a result, after co-evolution for 200 generations, the cooperators’ fitness is greater than that of the cheaters’, as shown in the payoff matrix in **Figure 1E**. This example illustrates how, in a duration of 200 generations, the payoff matrix associated with a cheater cooperator interaction can change quantitatively.

### Case II. Qualitative alterations to the structure of a payoff matrix – collapse of a cheater-cooperator game in melibiose

Melibiose is a glucose-galactose disaccharide, which is hydrolysed outside the cell by an α-galactosidase encoded by MEL1 (**Figure 2A**). Unlike the sucrose example, all the monosaccharides released after melibiose hydrolysis are free to be utilized by any individual in the population. Strains lacking MEL1 are unable to grow in media with melibiose as the sole carbon source^20^.

**Figure 2.**
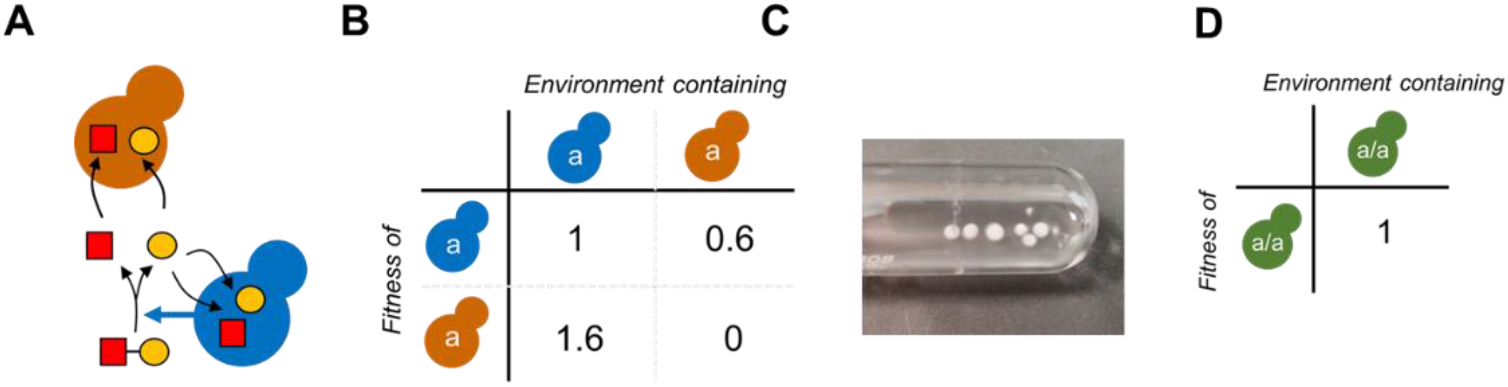
Collapse of a cheater-cooperator game in melibiose system. **(A)** Melibiose hydrolysis into glucose and galactose takes place in the extracellular environment due to release of MEL1-encoded α-galactosidase by the cooperator (blue). The resulting monosaccharides (glucose is represented by yellow circle and galactose by the red square) in the media and are available for anyone in the population. The cheater (orange) grows by utilizing the glucose and galactose thus released. **(B)** The payoff matrix of the system, when a small number of cheaters/cooperators are introduced in a population of cooperators/cheaters. Cheaters perform better than cooperators, when present in a population of cooperators. **(C)** Upon co-culturing, the two participating genotypes form flocs. The floc was mechanically sheared and the co-evolution experiment continued. After a few transfers (typically, 3-5), the culture media was taken over by growth of vegetative cells. **(D)** The payoff matrix associated with the evolved population is as shown. Clearly, a two-player game has collapsed into one with a single player.

Like in the sucrose case, we performed a similar co-evolution experiment with a strain carrying a functional MEL1 gene (cooperator) and a strain lacking MEL1 (cheater). As expected, the cheater, in the co-culture condition, exhibits a higher fitness than the cooperator, indicating a payoff matrix as shown in (**Figure 2B**).

When co-evolved, however, the two strains exhibit flocculation (**Figure 2C**). Upon shearing the flocs and plating for individual colonies, both cheater and cooperator were found to be present in the aggregate. After a few passages (∼4-6), the culture was taken over by a monoculture, which comprised of the hybrid between the two participating genotypes. The two distinct genotypes which we started the experiment with were lost from the culture.

This example provides an illustration of a scenario where two competing genotypes combine to create a fitter genotype, and in the process annihilate the two competing genotypes. In terms of the payoff matrix, the flocculation events lead to a two-player game being reduced to a one player game, as illustrated in the payoff matrix in **Figure 2D**.

### Case III. Qualitative alterations to the structure of a payoff matrix – birth of a cheater-cooperator game in melibiose

Melibiose utilization in *S. cerevisiae* is dictated by a peculiar regulatory network. Melibiose hydrolysis results in glucose and galactose. Between the two monosaccharides, glucose is the preferred sugar and is utilized before galactose. However, glucose represses expression of MEL1-encoded α-galactosidase. A pause in melibiose hydrolysis would imply that glucose in the media is exhausted, and the population then switches to galactose. However, utilization of galactose leads to MEL1 induction. The production and subsequent release of MEL1-encoded α-galactosidase re-initiates melibiose hydrolysis, leading to release of glucose (along with galactose) in the media (**Figure 3A**). In such a context, it is not clear what the metabolic commitment of the population will be at a single-cell resolution, when allowed to grow in melibiose.

**Figure 3.**
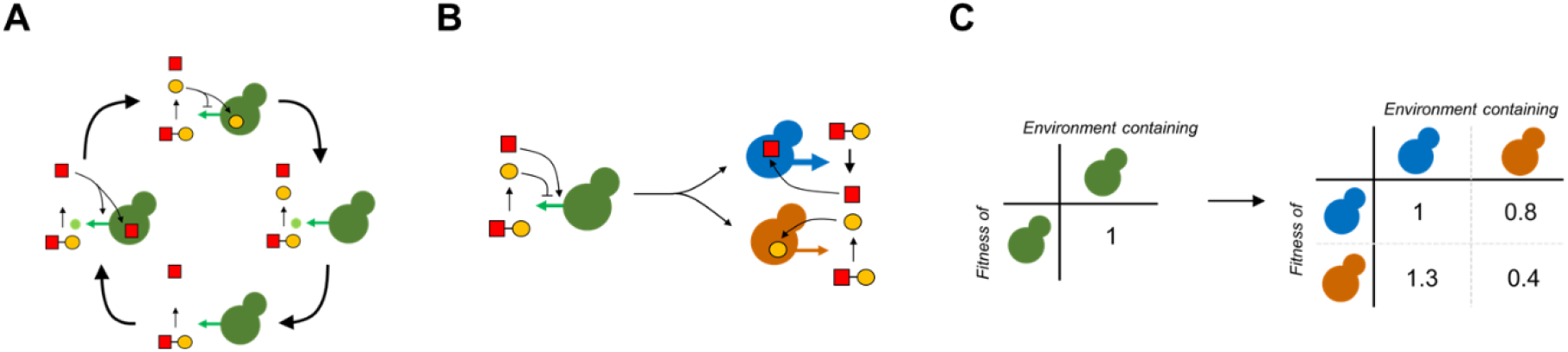
Emergence of a cheater-cooperator game after evolution of an isogenic population in melibiose. **(A)** Melibiose hydrolysis releases glucose (yellow circle) and galactose (red square). Of the two, glucose is preferentially utilized by *S. cerevisiae*. However, glucose utilization halts release of MEL1. Upon exhaustion of glucose, cells switch to galactose. Uptake of galactose also leads to induction of MEL1p production and release, resulting in melibiose hydrolysis and release of glucose. **(B)** After evolution for 400 generations, two genotypes were identified in the population. The cooperator (blue) acquired mutations in GAL3 locus of the GAL network, resulting in greater release of MEL1p and faster growth on galactose/melibiose. The cheater (orange) grew faster on glucose. **(C)** Starting with an isogenic population of *S. cerevisiae*, the population split into two distinct genotypes. This results in birth of a cheater-cooperator game.

Several possibilities exist. First, the population could switch from glucose to galactose and back. Such transitions, however, are detrimental to fitness^21^ and hence, such a strategy will likely not be successful. Second, mutations that abolish the hierarchy in glucose and galactose utilization could spread in the population, leading to a co-utilizer taking over the population. However, the nature and quantum of mutational events needed to make that transition to a co-utilizer is not well understood. Last, an isogenic population could split into two – each exhibiting a distinct state. In the case of melibiose, we could have a scenario where one fraction of the population produces MEL1-encoded α-galactosidase and utilizes galactose. The remaining fraction of the population grows on the glucose resulting from melibiose hydrolysis.

To test which of the three possibilities evolves, Mahilkar *et. al*.^22^ performed laboratory evolution with an isogenic population of *S. cerevisiae* in melibiose. After evolving for 300 generations, the isogenic population comprised of cells in two distinct phenotypic states. One part of the population expressed GAL1p, and the remaining fraction did not. After a few hundred generations of evolution, the population, genetically splits into two groups. Members of one of the two groups is a specialist at utilizing galactose, and the remaining fraction comprises of individuals less adept at utilizing galactose (**Figure 3B**). This split leads to two genotypes coexisting in the population. Therefore, a game with a single player has evolved into one with two players (**Figure 3C**).

## Discussion

Game theory was introduced to biology by John Maynard Smith and George Price 50 years ago^23^. Since their seminal work, game theory has found numerous applications in biology. A key assumption in evolutionary game theory is that participating players are rational agents^2^. In the context of microbial populations, this assumption is justified by assuming that the participating players are thought to have arrived at the optimal strategy in ecological time. However, populations are constantly evolving and accumulating genetic changes that help them improve their fitness^12,24^. Similarly, optimal strategies are also shown to change with feedback from the environment^25^. Therefore, changes in the payoff matrices are inevitable.

To study understand how payoff matrices evolve in microbial populations, we review and report simple laboratory evolution experiments with *S. cerevisiae* in two different sugar environments. We present empirical evidence for three types of changes that can occur in a payoff matrix of interacting microbial populations, after evolution for a few hundred generations.

While our results highlight the need to consider evolving payoff matrices while studying population dynamics, they also indicate how dramatic changes can occur in populations in short periods of time. In nature, environments are constantly changing and rarely ever consistent like in laboratory. Therefore, what are the strategies that wild populations use to avoid drastic changes in their structure? What is the optimal solution, in the face of constant environment fluctuations? Several studies have also pointed out how cooperation is maintained in populations that are spatially structured^26–28^. How then, do the payoff matrices evolve with change in spatial structure? Similarly, we do not yet know how our results would vary if the interacting pair of individuals belonged to different species. Therefore, our work motivates detailed high-throughput analyses directed towards building a comprehensive understanding of the evolution and applications of payoff matrices.

## Methods

### Strains used

BY4741 (*MATa his3Δ1 leu2Δ0 met15Δ0 ura3Δ0 SUC2*)^29^, a derivative of S288c, was used as the cooperator in Case I. SUC2 was replaced by the hygromycin resistance gene, as described in Raj and Saini^19^, to generate the cheater strain.

Haploid SC644 (*MATa MEL1ade1 ile trp1-HIII ura3-52*)^22^ was used as a cooperator and BY4741 was used as a cheater in Cases II and III.

### Calculation of payoff matrices

The cooperator and cheater strains were grown in allopatry in gly/lac media for 48 hours. To test the payoff of growth of strain *A* when grown in an environment comprising of strain *B*, the two strains were mixed in the environment of choice to a ratio 1:100 (*A*:*B*) to a starting OD of 0.1. The co-culture was allowed to grow to an OD 0.5, and the respective ratios of *A* and *B* quantified.

In the case of growth on sucrose (*Case I*), the ratio of cheaters to cooperators was estimated by plating the co-culture for single colonies on a YPD plate. The single colonies were replica plated on plate containing hygromycin to estimate the number of cheaters in the population. The growth rate *r* of the strain was estimated and normalized by the growth rate of the monoculture cooperator to give the payoff matrix element.

In the same manner, in case of melibiose (*Case II*), the ratio of cheaters to cooperators was estimated by plating the co-culture for single colonies on a YPD plate. The single colonies were replica plated on SCM glucose plates lacking either histidine (only cooperator will grow) or tryptophan (only cheater will grow). The growth rate *r* of each strain was estimated and normalized by the growth rate of the monoculture cooperator to give the payoff matrix elements.

In *Case II*, the flocs were verified by checking (a) for growth on the distinct auxotrophic markers on the two participating genotypes, (b) the DNA content of the resulting floc using FACS, and (c) for growth on melibiose.

